# Improving root characterisation for genomic prediction in cassava

**DOI:** 10.1101/695494

**Authors:** Bilan Omar Yonis, Dunia Pino del Carpio, Marnin Wolfe, Jean-Luc Jannink, Peter Kulakow, Ismail Rabbi

**Affiliations:** Montpellier SupAgro, 34060 Montpellier Cedex 02, France; Department of Plant Breeding and Genetics, Cornell University, Ithaca NY 14850; US Department of Agriculture - Agricultural Research Service (USDA-ARS), Ithaca, NY; International Institute for Tropical Agriculture (IITA), Ibadan, Nigeria; Department of Jobs, Precincts and Regions, AgriBio, Centre for AgriBioscience, Bundoora, Australia

**Author notes:** equal contribution (Bilan Omar Yonis, Dunia Pino del Carpio).

## Abstract

Cassava is widely cultivated due to its high drought tolerance and high carbohydrate-containing storage roots. The lack of uniformity and irregular shape of storage roots within and between genotypes poses significant constraints on harvesting and post-harvest processing. Routine assessment of storage root size and shape in breeding plots relies on visual scores.

Here, we phenotyped the Genetic gain and offspring (C1) populations from the International Institute of Tropical Agriculture (IITA) breeding program for root shape and size-related traits using image analysis of storage root photographs taken in the field.

In our study, using univariate genome-wide association analysis, we detected for most shape and size related traits, significant QTL regions located on chromosomes 1 and 12. The QTL region on chromosome 12 has been previously associated, using IITA breeding populations, to cassava mosaic disease (CMD) resistance.

Because the uniformity in size and shape of cassava roots is an important breeding goal, we calculated the standard deviation of individual root measurements per clone. The use of standard deviation measurements allowed the identification of new significant QTL for Perimeter, Feret and Aspect Ratio on chromosomes 6, 9 and 16. Using genomic prediction cross validation, the accuracies of root size and shape-related traits were lower than those previously reported for dry matter content (DM) and cassava mosaic virus resistance (CMD). Predictive accuracies of the mean values of root size and shape image-extracted traits were mostly higher than yield trait prediction accuracies in the C1 population. This study aimed to evaluate the feasibility of the image phenotyping protocol and to assess the use of genome-wide analyses for size and shape image-extracted traits. The methodology described here and the results obtained in this study are promising and open up the opportunity to apply high-throughput methods in cassava.

## Introduction

Cassava (*Manihot esculenta* Crantz), a tropical root crop with origins in Latin America, ranks as the 3rd most important crop in the tropics after rice and maize (Guira *et al*., 2017). In Africa, more than 800 million people rely on cassava as a primary source of calories (Howeler *et al*., 2013). Cassava is widely cultivated due to its high drought tolerance and high carbohydrate-containing storage roots, and although most of the production is for human consumption, its use extends to animal feed and industrially processed products (Hahn, Reynolds and Egbunike, 1992; Howeler *et al*., 2013; Lukuyu *et al*., 2014). In addition to the edible, high-starch storage roots, cassava plants produce thin fibrous roots, which function to absorb water and nutrients from the soil (Alves, 2002). The development and differentiation of fibrous roots, as well as the mechanism that triggers root storage formation in cassava, are poorly understood.

Cassava storage roots are morphologically diverse, the lack of uniformity and irregular shape between and within genotypes poses significant constraints on harvesting and post-harvest processing. The irregularity of root shape results in considerable losses of valuable root yield (Hahn, Reynolds and Egbunike, 1992). The waste of tuber flesh and the inefficiency of hand peeling could be avoided by peeling mechanization. However, breeding for root characteristics that facilitate this process requires a thorough understanding of the genetic basis of cassava root morphology. Several studies have attempted to characterize cassava root shape to support the development of peeling mechanization (Onwueme, 1978; Ejovo N. Ohwovoriole *et al*., 1988).The root characteristics that were evaluated in those studies include root diameter, weight, length and peel thickness.

Routine assessment of storage root size and shape in breeding plots relies on visual scores (www.cassavabase.org/search/traits). The categorical scores for root size are 3, 5 and 7 for small, medium and large roots, respectively. A single categorical score is given to a harvested plot based on the most frequent size in that plot. The visual rating of shape is 1 (conical), 2 (conical-cylindrical), 3 (cylindrical), 4 (fusiform), 5 (Irregular), and 6 (Combination of shapes). Similar to root size, the shape scoring is based on the most common observation in a plot. These categorical scores suffer from person to person subjectivity and inability to describe the variation in size and shape within a plot. Thus, image analysis of roots offers a more objective means of obtaining unbiased quantitative data on important root traits.

Image analysis software tools for high-throughput phenotyping have gained increased relevance due to the need in crop improvement to keep up with the advances in genotyping technologies (Furbank and Tester, 2011; Hartmann *et al*., 2011; Fahlgren, Gehan and Baxter, 2015). In Maize, imaging under controlled illumination followed by automatic image-analysis has been successfully used to study root system architecture traits (Colombi *et al*., 2015). In cereals, grain shape is an important target for genetic improvement, because it is usually related to quality, consumer appeal or the intended end usage (Lestrel, 2011). For rice grain shape description, SHAPE, a program based on Elliptical Fourier Descriptor (EFDs) has been used to derive shape-related phenotypes for genome-wide association and genomic prediction (Iwata *et al*., 2015b, 2015a).

Genomic selection (GS) is a method first introduced in animal breeding to select candidates for crossing in the breeding program using only genomic information. GS is particularly relevant for the improvement of polygenic traits (Heffner, Sorrells and Jannink, 2009) because its implementation can lead to a reduction in cost and time compared to traditional plant breeding programs (Jannink, Lorenz and Iwata, 2010). Because cassava is an outcrossing species mostly propagated by stem cuttings, conventional breeding methods can take more than five years to produce superior performing clones (www.nextgencassava.org). Genome-wide association studies (GWAS) are complementary to GS as they have proven effective for the identification of QTL regions associated with several traits that are critical for cassava breeding, including cassava mosaic disease resistance (CMD) (Wolfe *et al*., 2016), cassava brown streak disease resistance (CBSD) (Kayondo *et al*., 2018), and beta-carotene content and dry matter content (Rabbi *et al*., 2017).

In this study, size and shape related traits describing cassava roots were obtained through automated image analysis. We first estimated their heritability and conducted a genome-wide association study to explore the genetic architecture of cassava roots shape characteristics; then we compared the genomic prediction accuracy of image size and shape traits to those of root yield. Our research contributes to a better understanding of cassava root shape and explores the possibility of high-throughput phenotyping that would allow breeders to use GS to select varieties for quantitative root characteristics.

## Materials and methods

### Germplasm

We processed and analyzed cassava roots images taken from several field trials conducted by the International Institute for tropical agriculture (IITA) as part of their genomic selection breeding program. The cassava germplasm collections that we analyzed are known as Genetic Gain (GG) and the progeny of the first genomic selection event (C1), which are thus progeny of a subset of the GG population. The GG constitutes a large collection of important landraces, breeding lines and released improved varieties of cassava developed by IITA over the last four decades. More detail about the origins and constituency of these populations is available in several published studies (Wolfe *et al*., 2016;Wolfe *et al*., 2017).

A summary of the trials used in the present study is presented in Table 1. The first set of trial was the GG trial which comprised 805 plots planted in the summer of 2014 in Ubiaja, Nigeria using an augmented design with two checks planted in each incomplete block. The trial comprised of 758 unique clones. Each plot consisted of 10 stands in a single row with spacing of 1 m between rows and 0.8 m within rows. The second set of trials consisted of 88 clones selected from the GG population and planted as preliminary yield trial (PYT) across four locations (Ibadan, Ikenne, Ubiaja and Mokwa) using a randomized complete block design with two replicates. Plot size was similar to that of the GG trial. It is important to note here that these clones were used as parents for the GS cycle 1 population. The third set of trials involved GS cycle 1 clones that were split into three sets and planted separately in three locations: Ibadan, Ikenne and Mokwa. Each set was planted as a clonal evaluation trial (CET) using an incomplete block design with common checks in each block. All trials had at least 10 clones in common. Plants were harvested after 12 months in all trials.

**Table 1.**
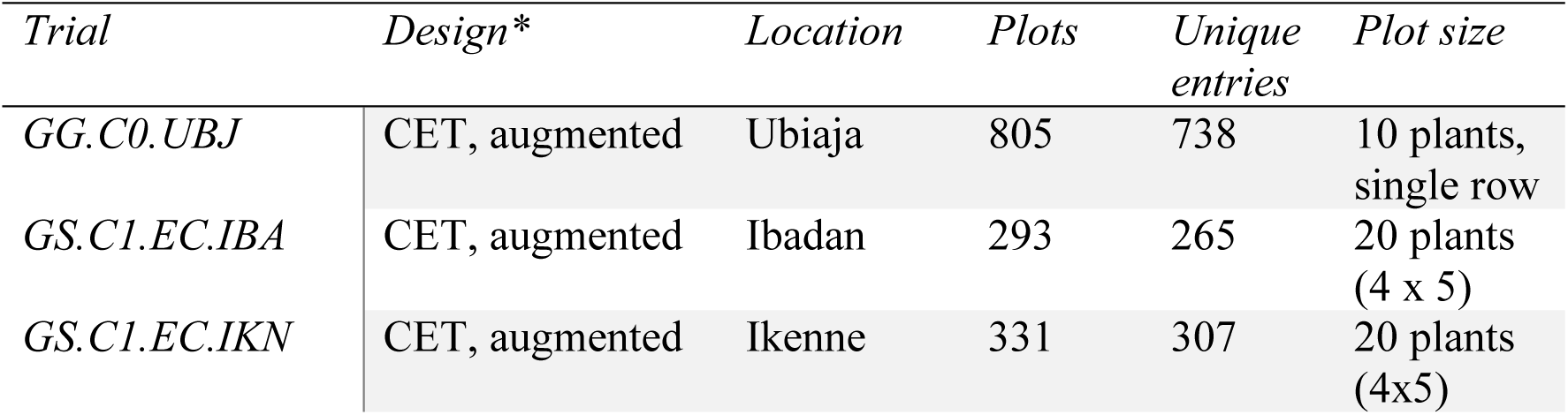
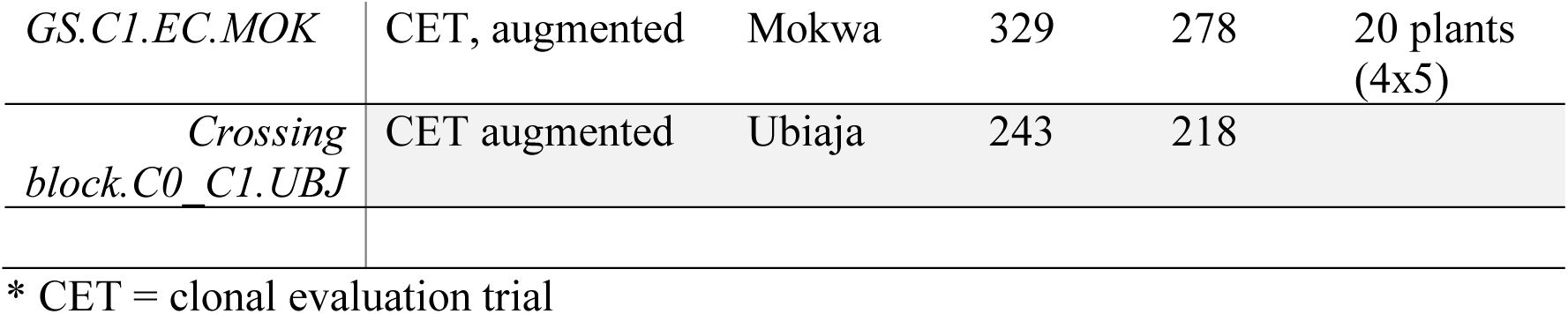
Summary of trials used in the present study including the trial names, design, locations, number of plots and number of unique clones in each trial.

### Image acquisition

The roots from four plants per plot were spread across a green board (160 cm by 120 cm). It was important that the roots were not touching each other and also not touching the board edges to get an individual root value (Supplementary Figure 1). Five circles, each 7.5 cm in diameter were painted on the left and right sides of the board. Those circles were used as a reference to transform the final result from the pixel unit to cm. Labels were placed on the board for each image allowing images to be identified and renamed for further processing.

### Image processing and phenotype acquisition

First, the images were coded to assign each photo to the plot from which the roots were taken. In some cases, several images were required per plot, to capture all roots from all the plants. For the GG collection, after quality control we obtained 805 images of cassava roots for 738 clones of which 665 had genotypic information. For the C1 population, we had images originating from four locations and a total of 1091 root images for 997 clones. All the image processing was performed with ImageJ Java version 1.8.0_11 (64-bit). The images were copied in two folders, one for processing and measuring the roots and the second for scaling the measurements. Thus, each image was processed and analysed twice.

#### Image processing

The first step of the image processing was to convert our RGB colour images into HSB stacks (hue, saturation and brightness images). We obtained three slices, but we only kept the first slice (the hue image). We then set a threshold from 0 to 255 for the roots and from 125 to 255 for the reference scaling circles before proceeding to run the “threshold” followed by the “make binary” commands. This threshold was determined by doing individual tests on some images. At the end of the processing, each image was binary, with our objects of interest (roots and scales) represented as white pixels and everything else as black. Most steps in the procedure were automated using customized ImageJ macros.

#### Phenotypes acquisition and description

The “analyze particles” command in ImageJ counts each contiguous area of white pixels within a binary image and gives some additional basic measurements. With the aim to get shape related traits, we used the “extended particle analyzer” function in the BioVoxxel Toolbox plugin (http://imagej.net/BioVoxxel_Toolbox#Extended_Particle_Analyzer). This function computes useful parameters of which we chose to keep seven for downstream analysis: Area, Perimeter, Feret, Circularity, Solidity, Roundness, and the Aspect Ratio (AR). The area and the perimeter describe the size of a root. The Feret, is the longest distance between any two points along the selection boundary, also known as maximum caliper. Circularity, Solidity, Roundness and aspect ratio (AR) describe shape.

The shape descriptors are ratio values that ranged from 0 to 1 except AR, which is not bounded. In addition, the shape descriptors do not have a unit, while area, perimeter, and feret are parameters expressed in pixels. The mean area value of the circles was used as a reference to convert pixels to centimetres (scaling coefficient). Since the exact diameter in centimetres of each circle was known, we used this value to calculate the mean number of pixels per cm^2^ for each image.

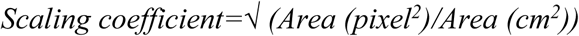

### Genomic analyses

We performed a two-step approach for the genomic analysis. In the first step, we used a linear mixed model to account for the variability in the field design and calculate the broad-sense heritability. The input data was: 1) the mean phenotype value for each plot (average phenotype of all imaged roots), 2) the same as (1) but adjusted to account for the potential effects of variation in cassava mosaic disease (CMD) severity among plots and 3) the standard deviation of the root shape and size measurements (across all imaged roots) per plot, also adjusted to remove the effect of CMD. We fit two different models, with CMD correction and without CMD correction, for each of the two focal populations (GG or C1).

For GG, the following models were fitted:

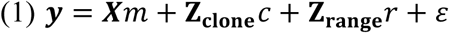

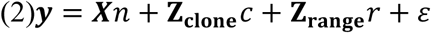

In both models ***y*** is a vector of phenotypes, **Z**_**clone**_ and **Z**_**range**_ are respectively the incidence matrices of the clones and range both fit as random with their effects vector 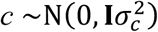 for clones and 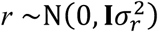 for range. **X** is the incidence matrix for the fixed effects. In model 1, the number of harvested plants per plot (NOHAV) and CMD were accounted for as fixed and the vector *m* contains the effect estimates. In model 2, we did not correct for CMD, **X** and *n* therefore only reference NOHAV.

For C1, model (3) and (4) were fitted:

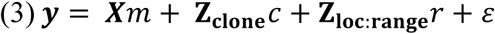

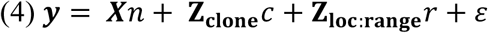

In model (3) and (4), we replace the range variable with the combination of the location and the range (Loc:range, i.e. range is nested in location) as the C1 population was planted in several locations unlike the GG. In all models, for all traits, ***y*** corresponded to the log transformation of the original phenotypic values. Additional explanation of the models fitted for these populations can be found in Wolfe et al. (2017).

From these models, we extracted the clone-effect BLUP, which estimates the total genetic value (EGV) of each line and de-regressed the EGV by dividing them by their reliability to obtain the de-regressed BLUP. Broad-sense heritability values were calculated using the variance components estimated using the mixed-models described above. EGV and de-regressed EGV are used in downstream analyses as described below.

### Genotyping data

Both populations (GG and C1) were genotyped using the genotyping-by-sequencing (GBS) method (Elshire *et al*., 2011). TASSEL 5.0 GBS pipeline v2 (Glaubitz *et al*., 2014) was used for SNP calling. Alignment of GBS reads was to the cassava reference genome v6.1 (http://phytozome.jgi.doe.gov; ICGMC, 2015). The condition for the genotype calls was the presence of a minimum of four reads. Extracted SNPs were filtered to remove clones with >80% missing and markers with >60% missing genotype calls. Markers were also removed when they had an extreme deviation from Hardy-Weinberg equilibrium (*χ*^2^ > 20).

A combination of custom scripts and common variant call file (VCF); manipulation tools were used to accomplish the above pipeline. The missing data were imputed using Beagle v4.0 (Browning and Browning, 2016). For the GG and C1 populations, we had 112,082 and 179,041 markers, respectively, with MAF > 0.01.

### Genomic prediction

We estimated genomic prediction accuracy using 5-fold cross-validation repeated 25 times similar to what is described in Wolfe et al. (2017). Briefly, the for each replicate of the process, the population was split into five approximately equal chunks (folds). Five genomic predictions were then made in which each fold (fifth of the population) in turn served as the test set (no phenotypes) and were predicted by the remaining four-fifths (training set, with phenotypes). Prediction accuracy for each fold was defined as the correlation of the genome-estimated breeding values (GEBVs, which are BLUPs from the test-sets of each fold), with the de-regressed EGVs from the pre-adjustment stage of the analysis.

For genomic prediction, we used a mixed-model with a genotype (clone) random effect with covariance proportional to the genomic relationship matrix, also called GBLUP. The genomic relationship matrix was constructed using the function *A*.*mat* in the R package rrBLUP (Endelman, 2011; Endelman and Jannink, 2012)De-regressed BLUPs were used as the response variable and the GBLUP models were fit with the function *emmreml* in the R package EMMREML (Akdemir and Okeke, 2015).

### GWAS analyses

Genome-wide association mapping (GWAS) analyses were performed using a linear mixed-model analysis (MLMA) implemented in GCTA (Version 1.90.0beta) (Yang *et al*., 2011). Specifically, we followed a leave-one-chromosome-out approach and tested all markers with MAF>0.05. The leave-one-chromosome-out approach involves excluding all markers on the chromosome of the current candidate SNP from the genomic relationship matrix (GRM) used to control population structure when estimating their marker effects. Manhattan plots were generated using the R package *qqman* (Turner, 2014) with a Bonferroni threshold of 6.28.

Candidate gene identification was performed using the significant GWAS results of the standard deviation + CMD correction GWAS results. Using the phytozome 12 portal link to biomart (https://phytozome.jgi.doe.gov/biomart/) we searched for genes located 10kb around the top SNP hits.

### Multivariate GWAS analysis

We used a multivariate linear mixed model as implemented in GEMMA (mvLMM) (Zhou and Stephens, 2014). We tested marker associations with multiple phenotypes that are fitted jointly in the mvLMM while controlling for population stratification. Different combinations of phenotypes were fitted in six models, the phenotypes that were fitted together were selected based on their phenotypic correlation. Model 1: Circularity, Round, Solidity; Model 2: Area, Feret, Circularity, Solidity, AR; Model 3: Area, Perimeter, Round, Solidity, AR; Model 4: Area, Perimeter, Feret, Circularity, Round, Solidity, AR; Model 5: Area, Perimeter, Feret; Model 6: Circularity, Round, Solidity, AR.

## Results

### Phenotypes distribution

Using the plugin BioVoxxel in ImageJ, we extracted quantitative measurements of the Area, Perimeter, Feret, Circularity, Solidity, Roundness, and the aspect ratio (AR) from root images collected in the field. The raw value datasets show similar ranges for root shape and size descriptors in GG and C1 populations (Supplementary Table 1). The individual root measurements with the maximum and minimum value of each trait in both populations are presented in Figure 1.

**Figure 1:**
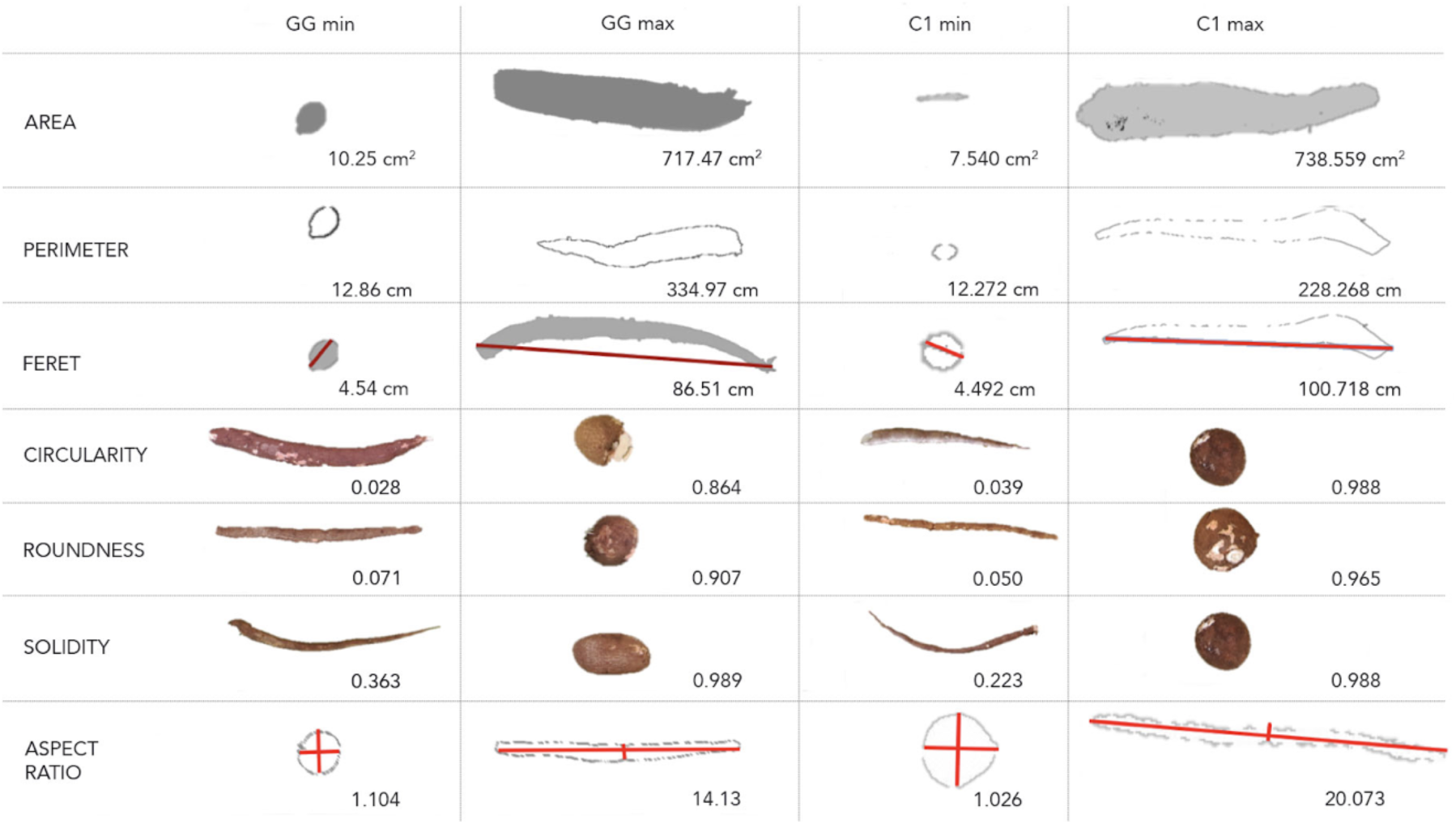
Phenotype description obtained using the extended particle analyzer plugin in ImageJ: Individual root measurements with the maximum and the minimum value of each trait in Genetic gain (GG) population and Cycle1 (C1) population, represented to highlight the range of values for each trait.

The frequency distribution of the mean value per plot of the GG and C1 populations is presented in Supplementary Figure 2 and the mean values per trait within population are presented in Supplementary Table 1. Some genotypes exhibited large differences in their mean values for Area, Perimeter and Feret. For example, the maximum mean root Area in GG population was 339 cm^2^ while the mean Area of the GG population was 121.5 cm^2^. Similarly, those genotypes exhibited a maximum mean root Perimeter of 135 cm and a maximum mean Feret of 50 cm while the mean Perimeter and Feret value in the GG population were 66 cm and 26 cm, respectively.

In the C1 dataset, the maximum mean value for the root area in the C1 dataset was 372 cm^2,^ while the mean area of that population was 128 cm^2^. The maximum values for Perimeter and Feret were 132 and 49 cm while the C1 population mean value for the two traits was 68 cm and 28 cm respectively.

### Correlation plots

Phenotypic correlations were calculated pairwise using de-regressed BLUPs of the mean values for each population separately (Figure 2). In the GG dataset, the highest correlation within yield traits corresponded to root number and root weight (r^2^ =0.79). Similarly, root number and root weight were highly correlated in C1 population (r^2^=0.88). In both datasets, correlations between yield traits were significant and high (r^2^ > 0.5) and these traits were also positively correlated with Area, Perimeter and Feret. However, a low correlation (r^2^ < 0.1) was observed between yield traits and root shape descriptors such as Circularity, Roundness, Solidity and AR in both populations.

**Figure 2:**
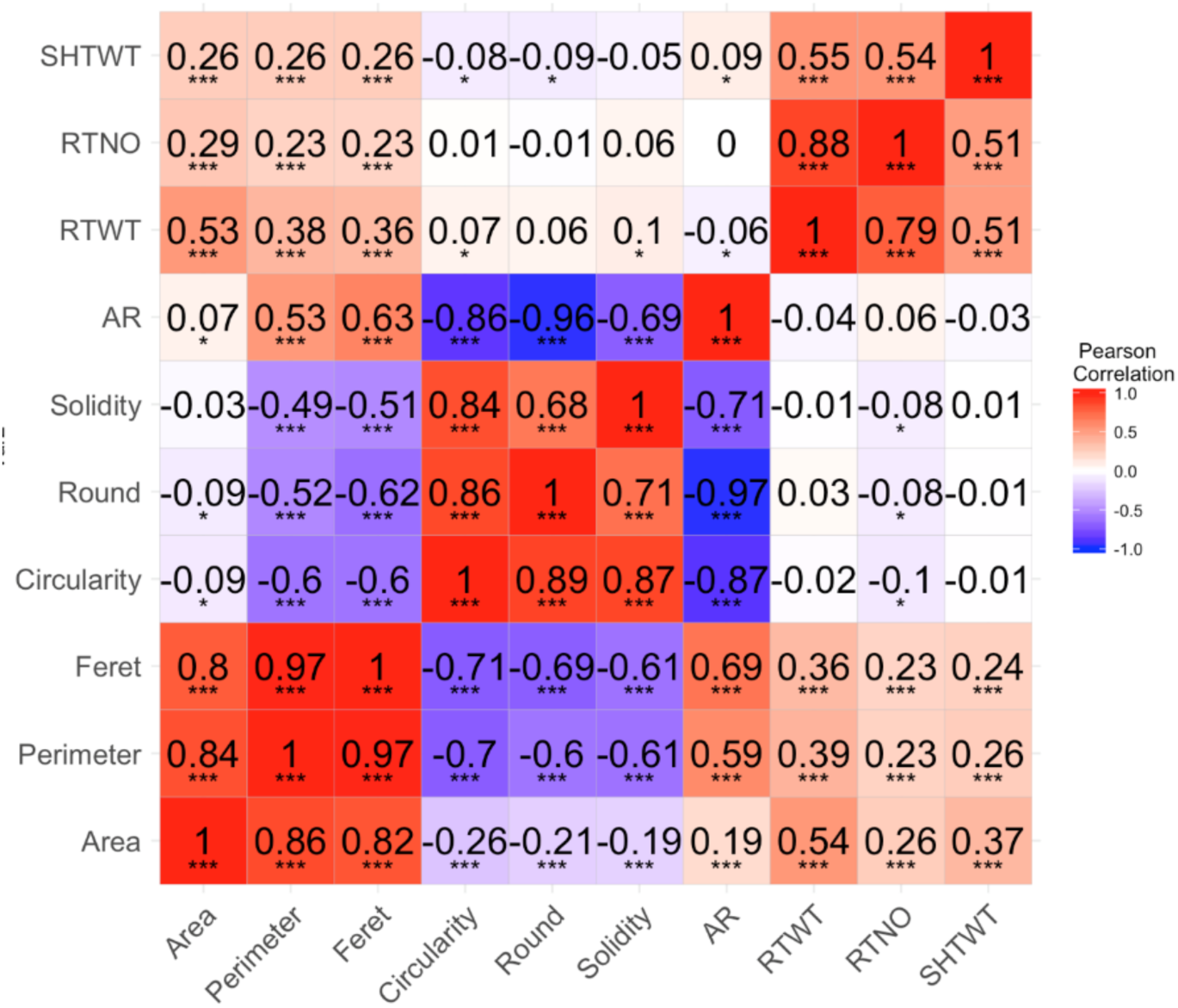
Heatmap with Pearson correlation coefficient: Trait correlation using the de-regressed BLUP value of GG dataset (lower triangle) and C1 dataset (upper triangle). The stars depict the significance according the p-value (\*\*\**P* < 0.0001, \*\**P* < 0.001, \**P* < 0.05)

Size-related traits derived from root images (Area, Perimeter and Feret) showed the highest positive correlation (r> 0.7) with each other. In both datasets, the highest correlation between size-related traits corresponded to Perimeter and Feret (r=0.97). Additionally, Feret and Perimeter were negatively correlated with shape-related traits (Circularity, Roundness and Solidity) and positively correlated with AR. In the GG dataset, Area showed a negative correlation with Circularity (r= −0.26), Roundness (r= −0.21), Solidity (r= −0.19), and a positive correlation with AR (r= 0.19). While in the C1 population, a low correlation was observed between Area and shape descriptors.

Within the shape related traits, the highest correlation was found between Circularity and Roundness (GG r = 0.89, C1 r = 0.86) and Solidity (GG r = 0.87, C1 r = 0.84). AR showed a negative correlation with Circularity, Solidity and Roundness in both datasets.

### Broad-sense heritability

Broad-sense heritability values (H^2^) for root shape and yield-related traits were calculated for each population (Table 2). In the GG population, without adjusting the phenotypes for their CMD score, H^2^ of root shape related traits ranged from 0.17 (Perimeter and Circularity) to 0.46 (aspect ratio) and for yield traits, H^2^ ranged from 0.29 root weight (RTWT) to 0.44 shoot weight (SHTWT). In the GG dataset, Perimeter, Circularity and Solidity exhibited the lowest heritability values at 0.17, 0.17 and 0.12, respectively.

**Table 2:**
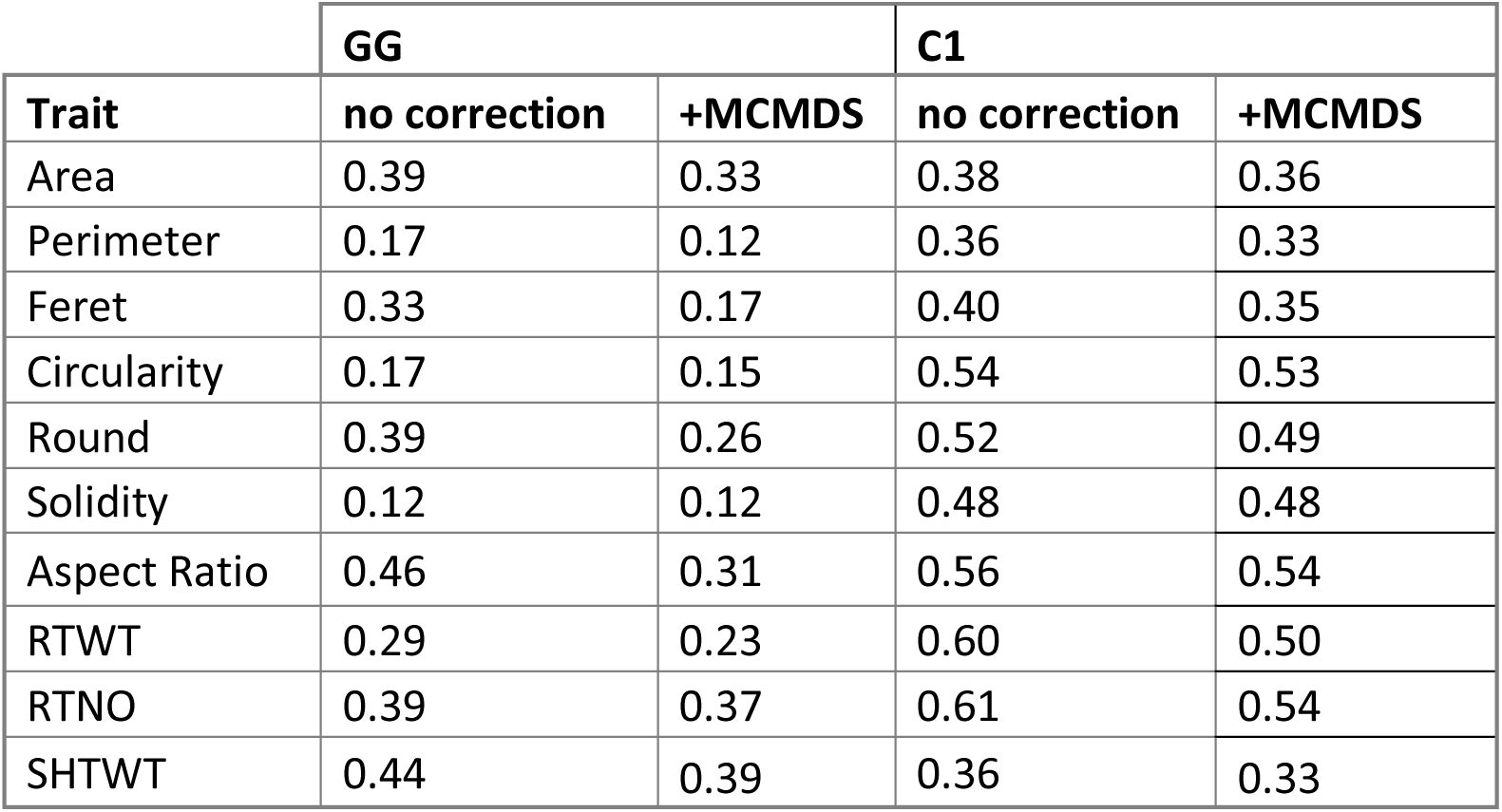
Broad-sense heritability values of root shape and root yield traits for the Genetic gain and cycle 1 breeding populations. (RTWT: root weight; RTNO: root number; SHTWT: Shoot weight).

In the C1 population, the heritability of shape-related traits ranged from 0.36 (Perimeter) to 0.54 (Circularity) while for yield traits H^2^ ranged from 0.36 (SHTWT) to 0.61 (RTWT). The heritability of most traits was higher in the C1 population than GG except for Area (0.39 to 0.38) and SHTWT (0.44 to 0.36). The inclusion of the CMD in the calculation of the variance components always reduced the heritability of all the traits in both populations by around 10%.

### Genome-wide association study of root traits

Using a univariate genome-wide association approach for root image traits (root size and shape) and root yield traits we identified significant loci for all traits except for area (Figure 3). We detected a total of 91 SNP markers exceeding the significance threshold (−log10 *P* ≥ 6.28). The Manhattan plots of the univariate GWAS results for yield traits are shown in Supplementary Figure 3 and detailed information on the significant markers is summarized in Supplementary Table 2.

**Figure 3.**
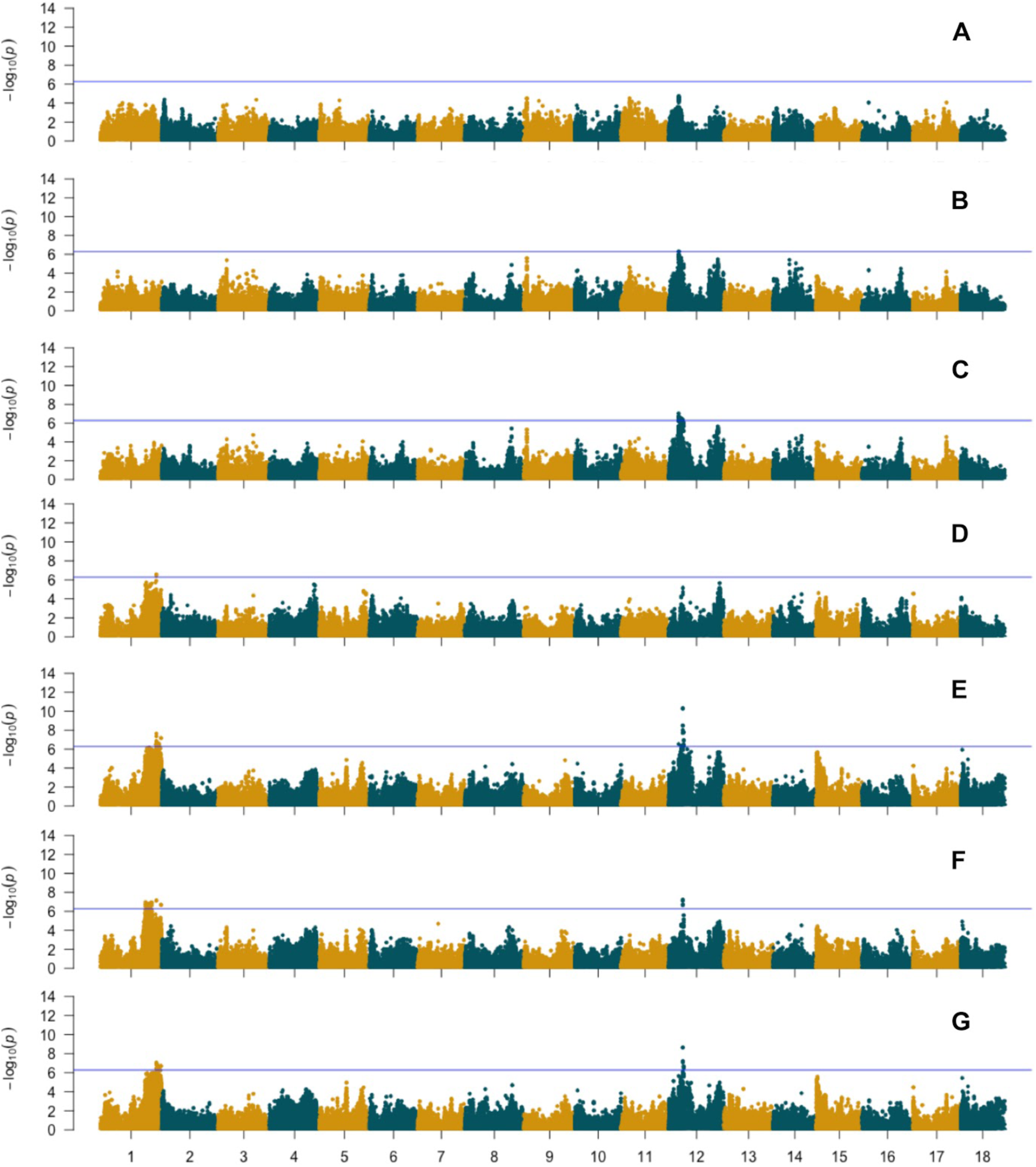
Genome-wide association results of size and shape-related traits using de-regressed BLUPs of mean values (not corrected for CMD). A. Area; B. Perimeter; C. Feret; D. Solidity; E. Aspect ratio; F. Circularity; G. Roundness. Blue horizontal line indicates the Bonferroni statistical threshold.

We detected markers associated with Perimeter and Feret on chromosome 12, and with Solidity on chromosome 1, whereas for AR we identified significant loci on chromosome 1 and chromosome 12. Similarly, for Circularity and Roundness, we detected significant loci on chromosome 1 and chromosome 12.

For most shape-related traits several other regions on chromosomes 3, 4, 8, 9, 14, 15 and 18 did not reach the significance threshold but showed a −log10 *P* ≥ 5 (Figure 3). For root yield traits we detected a QTL on chromosome 12 associated to root number (RTNO) and RTWT (Supplementary Figure 3, Supplementary Table 2). Notably, using the CMD adjusted phenotype removed the significance of the QTL on chromosome 12 but did not identify new QTL for the image traits shape phenotypes (Supplementary Figure 4) it detected new loci associated with root number and shoot weight (Supplementary Table 3).

Significant SNP markers ((−log10 *P* ≥ 6.28) were detected for the standard deviation-derived traits of Perimeter (per-sd), Feret (feret-sd) and Aspect Ratio (AR-sd) (Figure 4). For per-sd, a significant QTL was detected on chromosome 16, though it was not observed in the GWAS model with the mean values nor in the GWAS model with mean values with CMD adjusted phenotypes. For feret-sd, two significant QTL were identified, one on chromosome 9 and one on chromosome 6 and for AR-sd one significant QTL was found on chromosome 8 (Supplementary Table 4).

**Figure 4.**
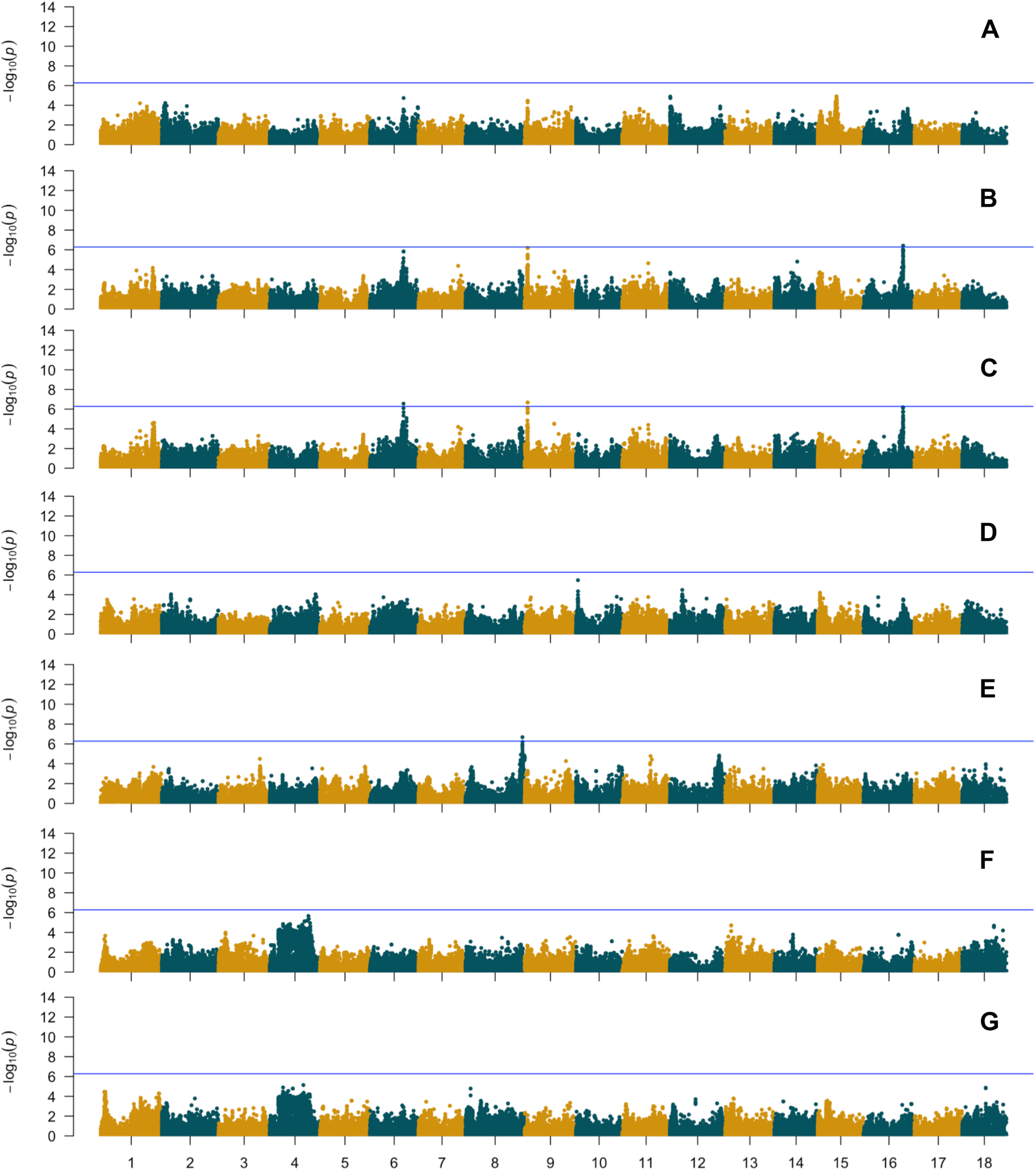
Genome-wide association results of standard deviation-derived size and shape-related traits using de-regressed BLUPs of mean CMD-corrected values. A. Area; B. Perimeter; C. Feret; D. Solidity; E. Aspect ratio; F. Circularity; G. Roundness. Blue horizontal line indicates the Bonferroni statistical threshold.

Different markers were significant in the multivariate GWAS model dependent on which phenotypes were included in the multivariate linear mixed model (mvLMM). Although the multivariate model can increase the power for detecting pleiotropic variants when using correlated traits, we identified few significant markers above the Bonferroni threshold (Supplementary Fig 5-10). Nonetheless, when P-values were corrected for multiple testing by computing Benjamini-Hochberg q-values, four SNPs were identified as significant in the multivariate analysis. In the multivariate analysis using Area, Perimeter, Feret, Circularity, Round, Solidity and AR in the mvLMM (Model 4) we identified a significant marker at the same location on chromosome 4 (Supplementary figure 8). Similarly, using model 6 (Circularity, Round, Solidity, Aspect ratio) we identified one significant marker located on chromosome 4 (Supplementary figure 10). When Area, Perimeter and Feret were included in the mvLMM (model 5) we identified significant markers on chromosomes 6 and 9 using a q-value threshold of < 0.1 (Supplementary figure 9).

### Genomic prediction

Using the parental (GG) and offspring generation (C1) datasets independently, we calculated the prediction accuracies of size and shape image traits and compared those to root yield traits accuracies using de-regressed BLUPs of 1) the mean phenotype value (average phenotype of 4 plants) (Figure 5, Supplementary table 5), 2) the mean root size and shape phenotypes adjusted to account for the potential effect of cassava mosaic disease (CMD) on these traits (Figure 6, Supplementary Table 5) and 3) the standard deviation of the root shape and size measurements adjusted to remove the effect of CMD (Figure 6, Supplementary Table 5).

**Figure 5:**
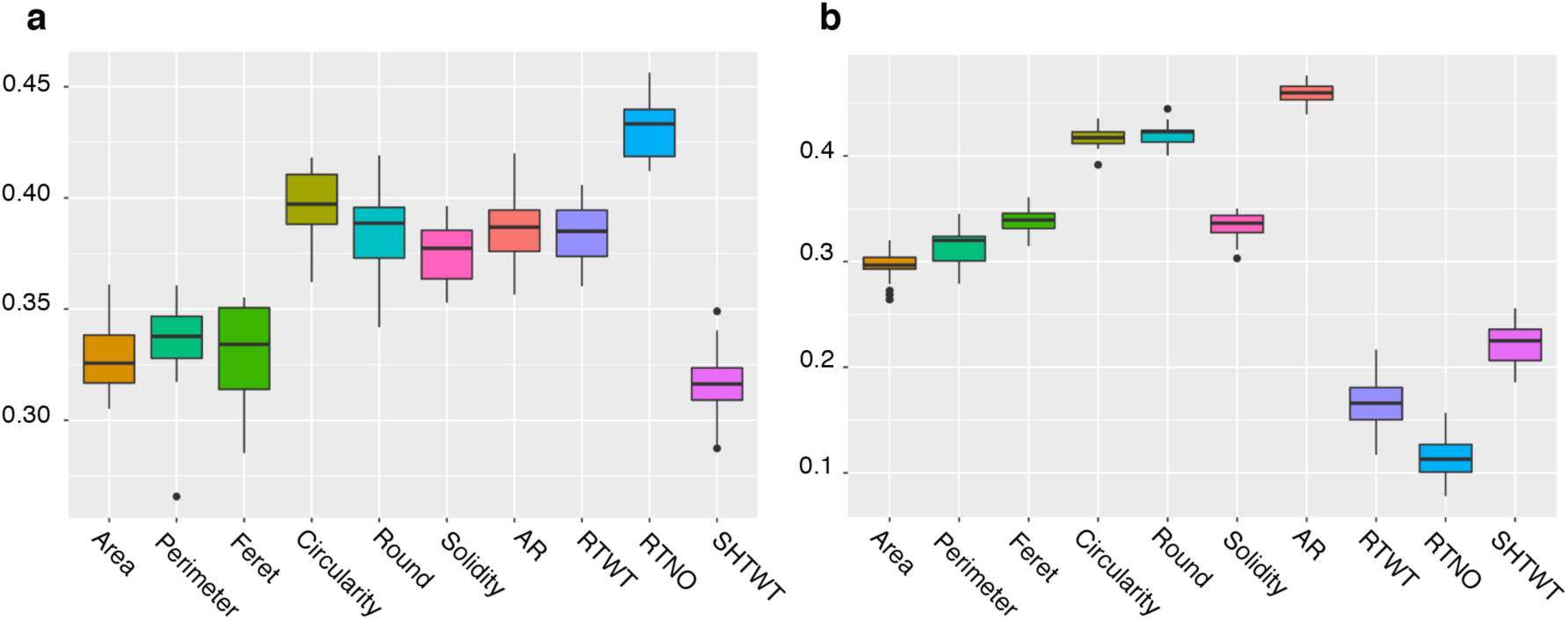
Prediction accuracy of root size and shape and yield traits. Predictive accuracies were obtained with 5 fold-cross-validation analysis using a GBLUP model in the (a) GG dataset and in the (b) C1 dataset.

**Figure 6.**
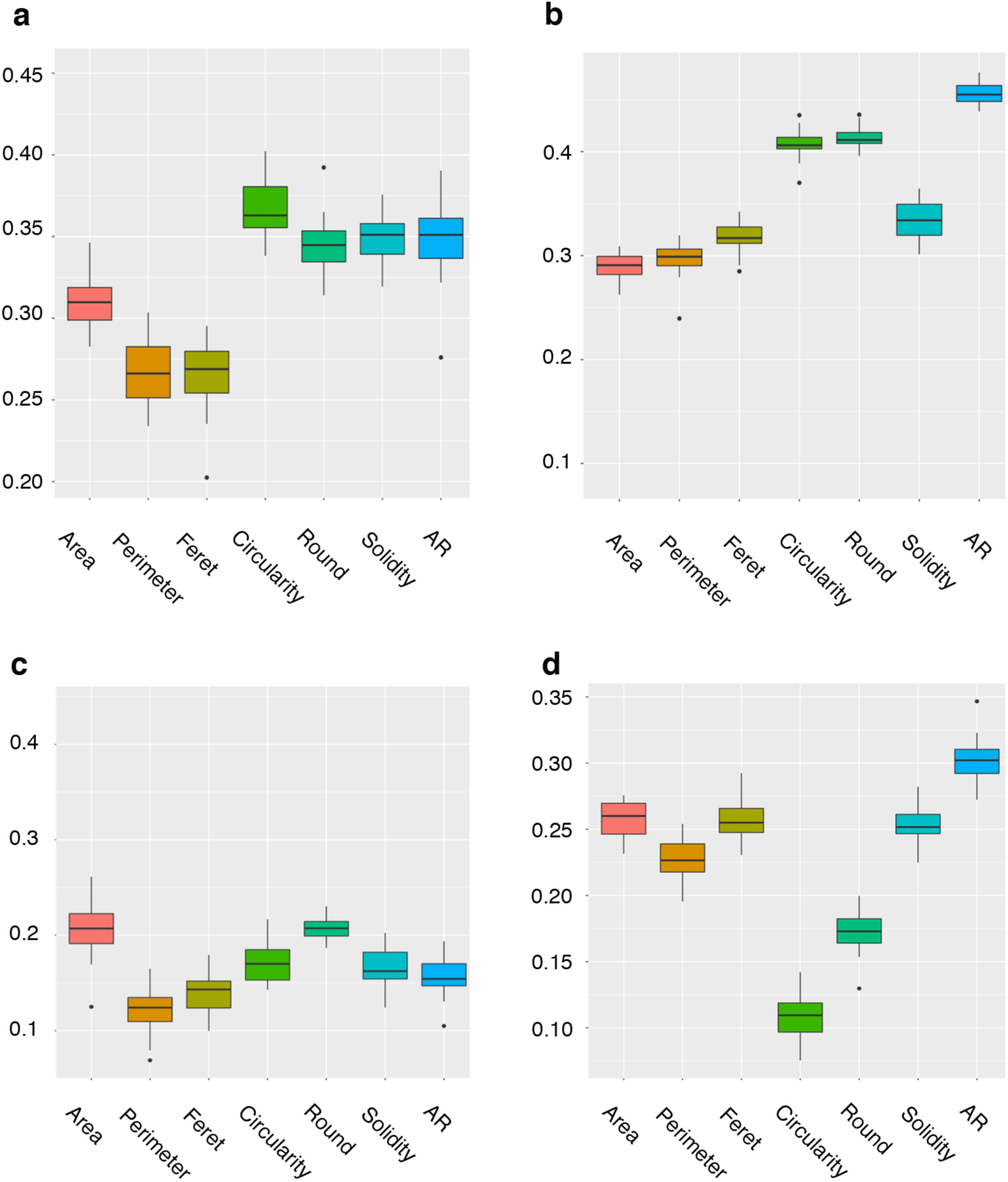
GBLUP model predictive accuracy of root size and shape traits. a) GG population CMD adjusted phenotypes, b) C1 population CMD adjusted phenotypes, c) GG population standard deviation + CMD correction, d) C1 population standard deviation + CMD correction

Prediction accuracy, calculated as the correlation between the genome estimated breeding values (GEBVs) and the de-regressed BLUPs of the mean phenotype value, ranged from 0.32 (SHTWT) to 0.43 (RTNO) in the GG population and from 0.12 (RTNO) to 0.46 (AR) in the C1. For yield traits, accuracies in GG were higher than in C1 but were not different between populations for the shape and size related traits. In the GG population, the shape descriptors Circularity (mean = 0.40), Roundness (mean=0.39), Solidity (mean =0.37) and AR (mean =0.38) showed slightly higher accuracies than the size descriptors Area (mean =0.33), Perimeter (mean = 0.34) and Feret (mean = 0.33). In the C1 population, size and shape image traits exhibited a higher prediction accuracy than root yield traits. Among the size descriptors, Feret showed the highest accuracy (mean=0.34) and Area the lowest (mean=0.29). Among shape descriptors, AR showed the highest predictive value (mean=0.46) and Solidity the lowest (mean=0.33) (Supplementary Table 5). When the mean root size and shape phenotypes were adjusted to account for the effect of CMD, we observed a minimal decrease in predictive accuracy (Supplementary Table 5). A lower predictive accuracy was obtained for standard deviation of size and shape traits adjusted for CMD, in both populations. In the GG population, the decrease was pronounced with a maximum reduction of up to 55% for root perimeter (0.27 mean to 0.12 CMD adjusted) while in the C1 population the largest reduction was of 73% for circularity (0.41 mean to 0.11 CMD adjusted) (Supplementary Table 5).

## Discussion

Root number and root weight are among the most important targets for improvement in cassava breeding programs. Although cassava root characterisation has been the subject of several studies (Adetan, Adekoya and Aluko, 2003; Padonou, Mestres and Nago, 2005; Anggraini *et al*., 2009), the genetic architecture underlying cassava root shape remains unexplored. This study aimed to evaluate the feasibility of the image phenotyping protocol and to assess the use of genome-wide analyses for size and shape image-extracted traits.

Here, we phenotyped the GG and C1 populations from the International Institute of Tropical Agriculture (IITA) breeding program for root shape and size-related traits using image analysis of storage root photographs taken in the field. In both populations, the storage roots exhibited a wide range of shape variation. Root-size related traits (Area, Perimeter and Feret) obtained through image analysis showed significant but low correlation (r ≤ 0.5) with cassava root yield components (RTNO, RTWT). Roots with a large area were generally heavier and the circularity of storage roots was mostly inversely correlated to its area. These results, suggest that rounded-shaped roots in cassava are generally smaller and hence lighter in weight. More importantly, the lack of correlation between size-related traits and shape related traits increases the interest in shape related traits as a target for selection.

In radish, rice and wheat, imaging-based studies of root shape and size traits have demonstrated first, that these have different genetic architectures (Iwata *et al*., 2000) and second, that shape phenotyping can aid the identification of pleiotropic QTL. In our study, using univariate genome-wide association analysis, we detected for most shape and size related traits, significant QTL regions located on chromosomes 1 and 12. The QTL region on chromosome 1 has been previously shown to be segregating for an introgressed segment from *M. glaziovii* (Bredeson *et al*., 2016). Furthermore, the QTL region on chromosome 1 has been associated, in the IITA genetic gain population, with other root traits such as dry matter and total carotenoid content (Rabbi *et al*., 2017).

For root weight and root number, we identified a significant QTL associated with those traits on chromosome 12. The QTL region on chromosome 12 has been previously associated, using IITA breeding populations, to cassava mosaic disease (CMD) resistance (Wolfe *et al*., 2016). The effect of cassava mosaic disease (CMD) on root yield has been previously investigated in fully and partly infected stands of cassava (Seif, 1982; Otim-Nape, Thresh and Shaw, 1997; Owor *et al*., 2004). In those studies, fresh stem, leaf and root yields and the number of tuberous roots were influenced by the health status of the plants harvested and that of their nearest neighbours. In our study, when we adjusted the size and shape phenotypes according to their CMD score we did not identify new QTL but a reduction in marker significance, which suggest that the CMD2 locus in chromosome 12 does not participate in the regulation of size and shape phenotypes. Nonetheless, the identification of new QTL for root number and shoot weight, when these traits were adjusted according to the CMD score, support the notion that CMD can have an effect on root yield traits.

Because the uniformity in size and shape of cassava roots is an important breeding goal we calculated the standard deviation of individual root measurements per clone. The use of standard deviation measurements allowed the identification of new significant QTL for Perimeter, Feret and Aspect Ratio on chromosomes 6, 9 and 16. For the new QTL regions located on chromosomes 9 and 16 we identified candidate genes related to the tocopherol and carotenoids pathways which are known regulators of plant development (Nisar *et al*., 2015) (Supplementary table 6). On chromosome 6, the most promising candidate is Manes.06G078700 a root meristem growth factor 1 related gene.

Together our GWAS results suggest that 1) root-related traits have in common the genetic control under few large effect loci and many small effect loci, 2) a possible correlation between disease severity and yield loss and, 3) that introgressed regions contain gene clusters which control root yield and root size/shape traits.

To increase the power of our study and to detect pleiotropic loci for size and shape traits (Korol *et al*., 2001; Korte *et al*., 2012), we used a multivariate linear mixed model approach which included groups of correlated root size and root size/shape traits. Considering multiple phenotypes in the mvLMM enabled us to identify new candidate loci on chromosomes 4, 6 and 9 that were not identified in the univariate analyses.

The potential of GS as a breeding tool to increase the rates of genetic gain was recently tested in three Next Generation Cassava Breeding programs (Marnin D. Wolfe *et al*., 2017). The study showed promising results particularly for traits with consistent heritability values across programs and stable large-effect quantitative trait loci. Prediction accuracies for RTNO, RTWT and SHTWT were similar with those reported in previous cassava cross-validation analyses (Wolfe *et al*., 2017). Root size and shape-related trait accuracies were lower than those reported for dry matter content (DM) and cassava mosaic virus resistance (CMD) (Wolfe *et al*., 2017). Although the heritability of yield traits was higher in the offspring (Cycle 1, C1) than the parental generation (Genetic Gain, GG), the predictive accuracy of traits extracted from root images showed intermediate to high values in both populations. However, the C1 yield traits accuracies being lower than the GG, suggests that because the C1 had been selected strongly for these yield traits, its variance was diminished.

Nonetheless, predictive accuracies of the mean values of root size and shape image-extracted traits were mostly higher than yield trait prediction accuracies in the C1 population. Adjusting the mean and standard deviation phenotypes for the effect of CMD reduced the predictive accuracy. However, that correction is necessary to unlink the effect of CMD from the causal loci that are responsible for the regulation size and shape root traits.

Although these measurements were laborious in the field and not high-throughput, the analyses of the images are automated and quantitative, they avoid subjectivity in scoring and other human-errors and most importantly, they improve cassava root characterisation. The methodology described here and the results obtained in this study are promising and open up the opportunity to apply high-throughput methods in cassava. The image capture and analysis can now be performed using the OneKK (one thousand kernels) app (https://github.com/PhenoApps/OneKK), an inexpensive and user-friendly tool for automated measurement of seed size, shape, and weight using smart phones. The app is developed under the BREAD PhenoApps project and supported by the National Science Foundation. Still, there is a need to explore the use of image-based phenotyping in multiple environments to estimate the effect of the environment on root shape related traits and to automate the collection of root images in the field further.

## Acknowledgements

This research was supported by the Bill & Melinda Gates Foundation and the Department for International Development of the United Kingdom through the “Next Generation Cassava Breeding Project”, and CGIAR-Research Program on Roots, Tubers and Bananas. The authors acknowledge the support of Andrew Smith Ikpan, Ogunpaimo Kayode and Cynthia Idhigu in acquiring the root images. We also acknowledge the PhenoApp project led by Jesse Poland, Kansas State University, for technical support in image-based phenotyping using their OneKK (one thousand kernel) application.

## Supplementary figures

**Supplementary figure 1:**
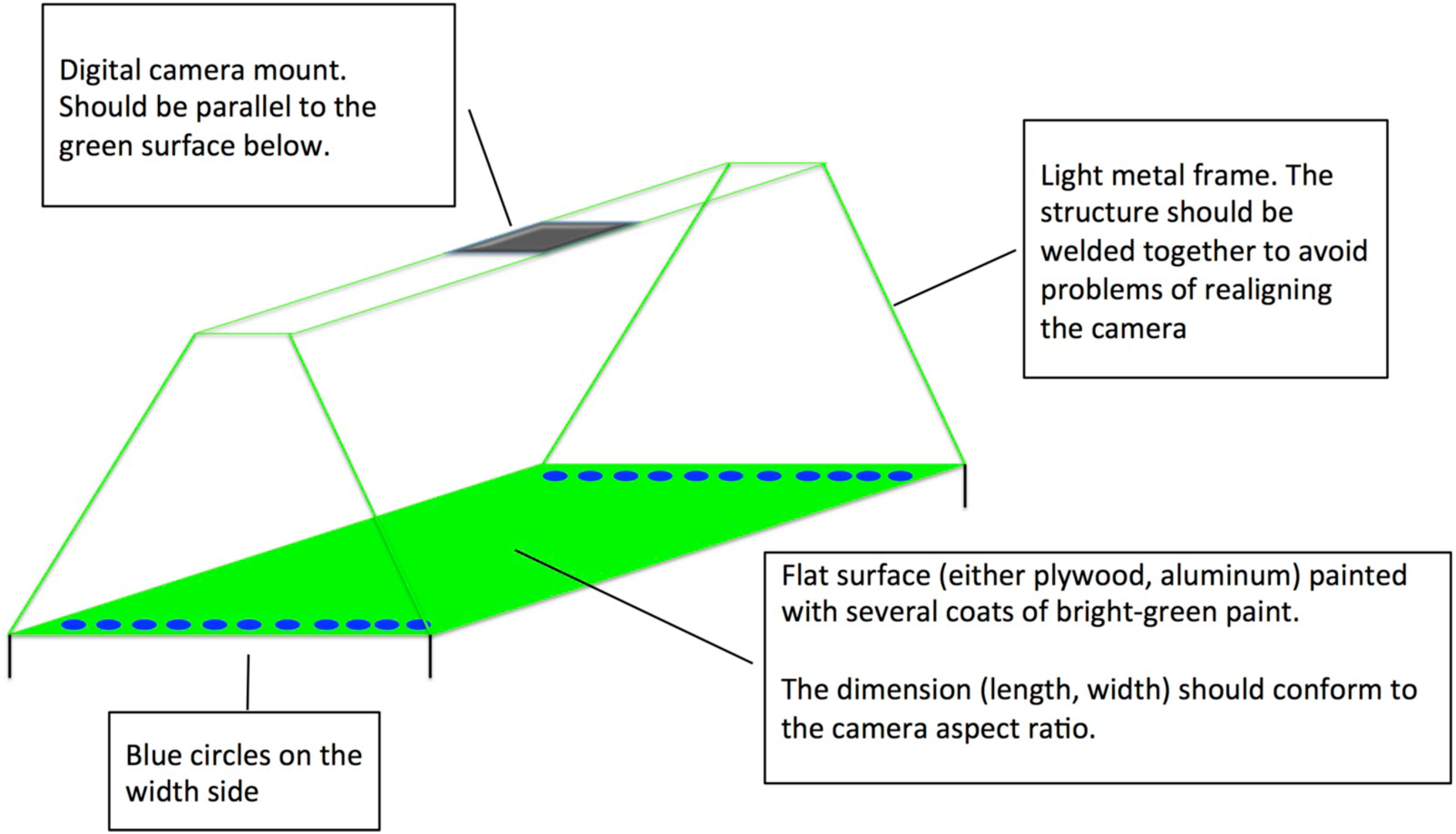
Schematic of the green board used as a background to take photographs in the field.

**Supplementary Figure 2.**
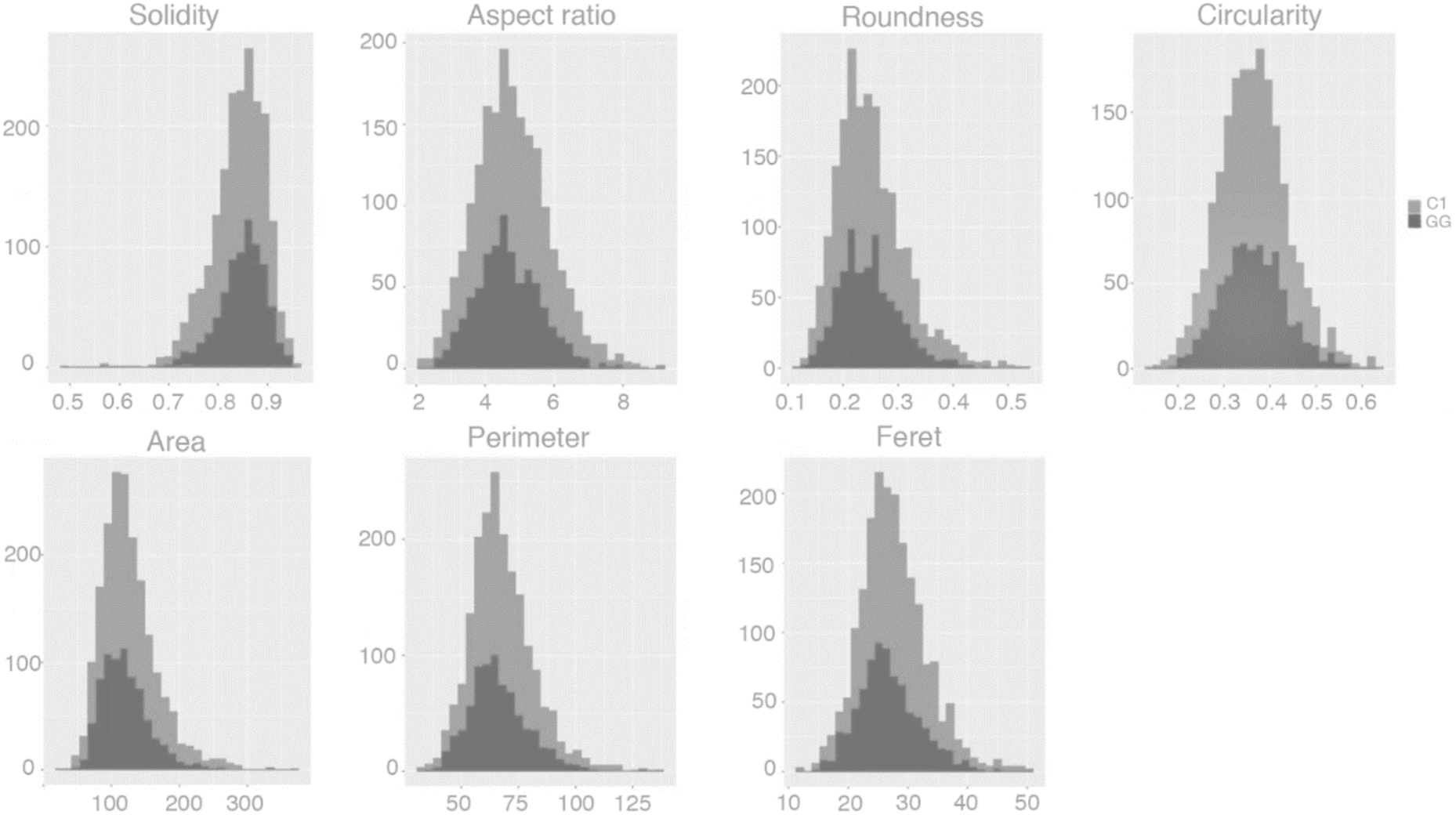
Trait distribution of mean root values. Root values of each phenotype extracted from photographs from each genotype were averaged and the distribution plotted. Dark gray: GG, light gray : C1.

**Supplementary figure 3.**
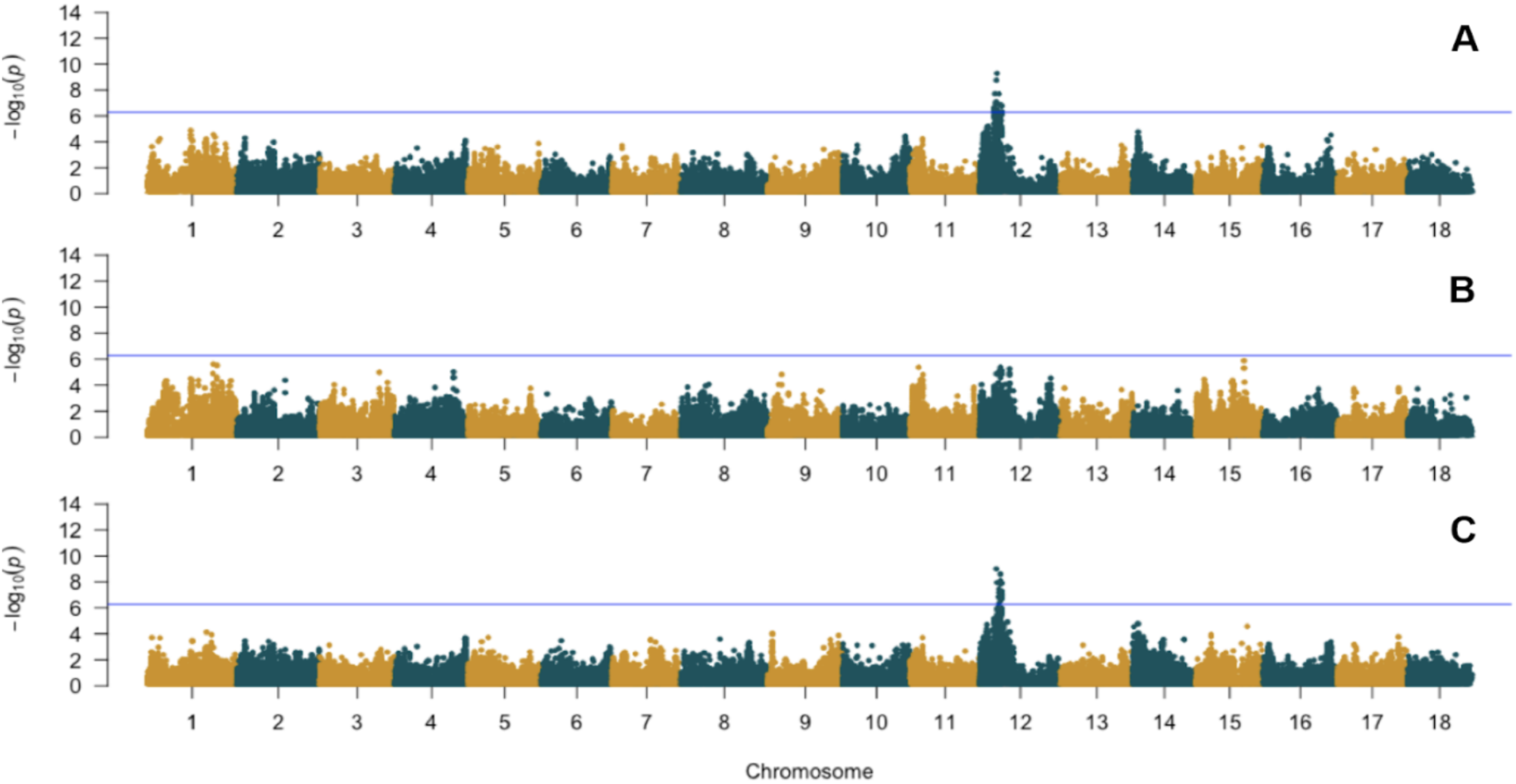
Yield traits GWAS results using the IITA Genetic Gain and Cycle 1 breeding populations. Manhattan plots of −log_10_(P-value) of A. Root weight, B. Shoot weight, C. Root number. Blue horizontal line indicate the Bonferroni statistical threshold (−log_10_(P-value) >6.28)

**Supplementary figure 4.**
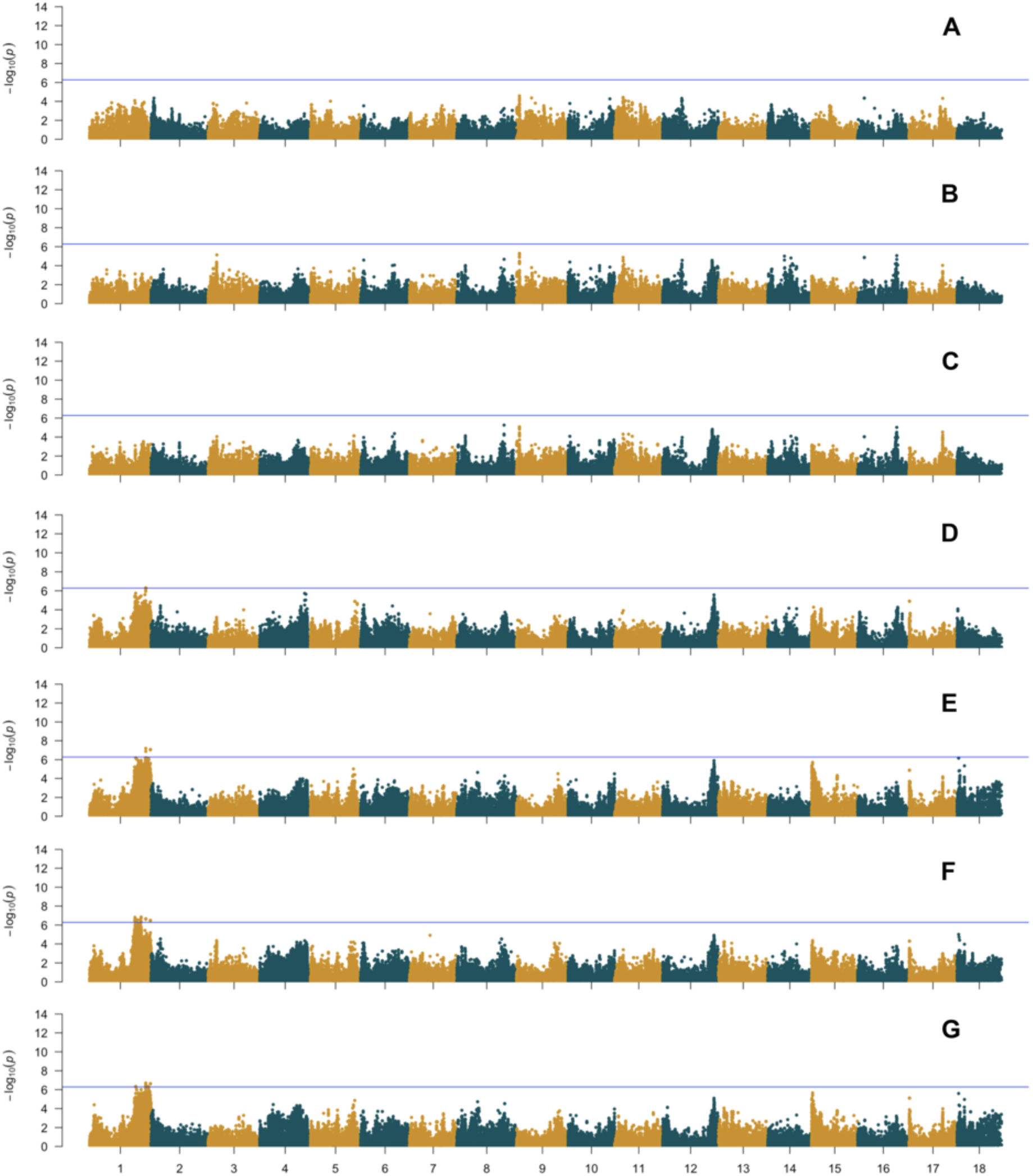
Size and shape GWAS results using the IITA Genetic Gain and Cycle 1 breeding populations using CMD score as a covariate. Manhattan plots of −log_10_(P-value) of A. Area; B. Perimeter; C. Feret; D.Solidity; E. Aspect ratio; F. Circularity; G. Roundness. Blue horizontal line indicates the Bonferroni statistical threshold.

**Supplementary Figure 5.**
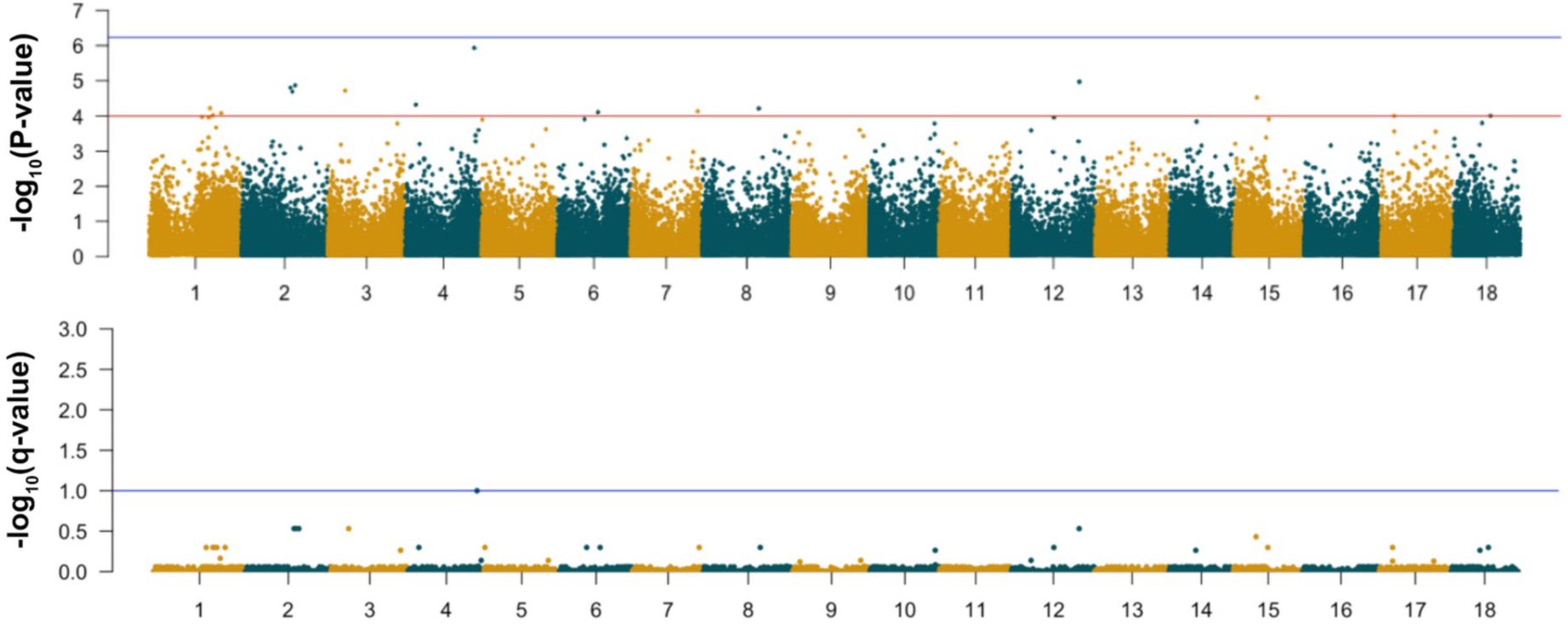
Multivariate GWAS of Circularity, Round and Solidity (model 1). Manhattan plots of −log_10_(P-value) (top panel) and −log_10_(q-value) (bottom panel). In the top panel the blue horizontal line indicates the Bonferroni statistical threshold and the red line indicate a −log_10_(p-value) = 4. In the bottom panel de the blue line indicates the significant threshold of the q-value.

**Supplementary Figure 6.**
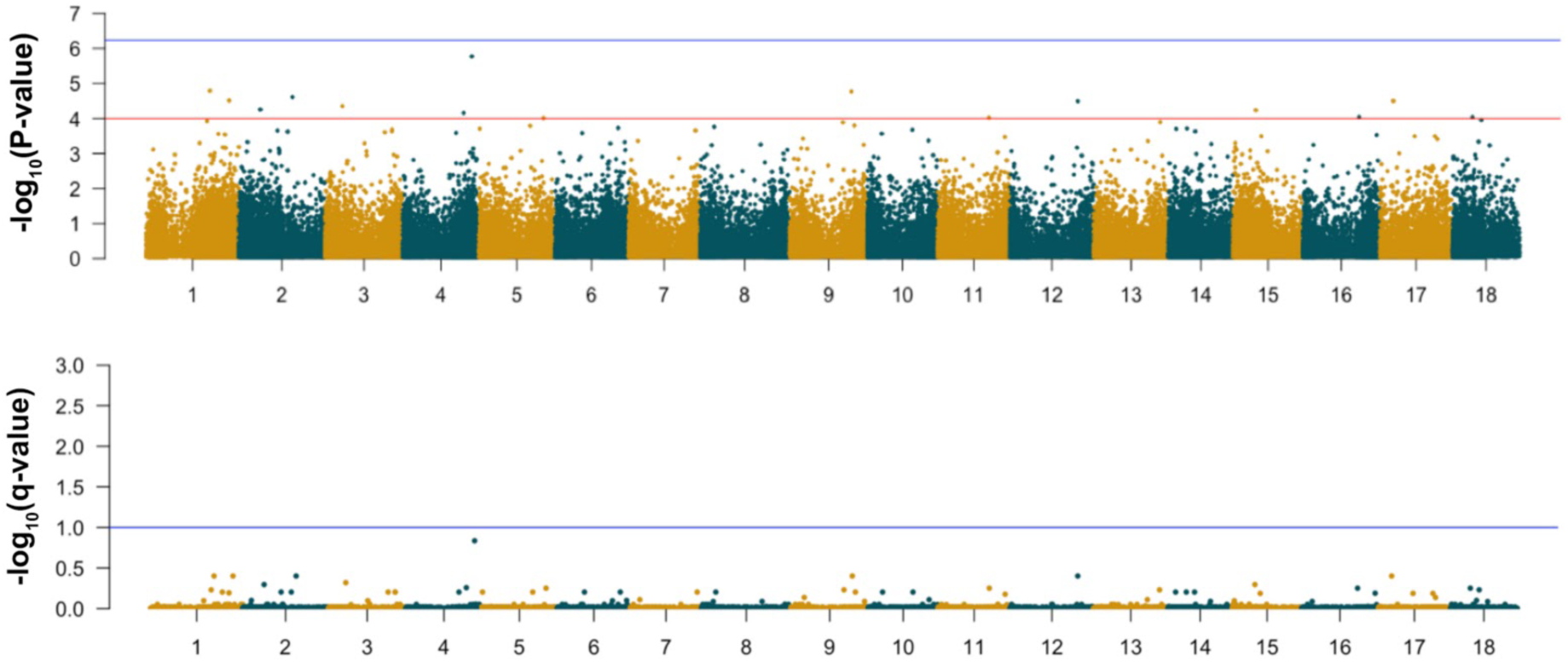
Multivariate GWAS of Area, Feret and Circularity (model 2). Manhattan plots of −log_10_(P-value) (top panel) and −log_10_(q-value) (bottom panel). In the top panel the blue horizontal line indicates the Bonferroni statistical threshold and the red line indicate a −log_10_(p-value) = 4.In the bottom panel de the blue line indicates the significant threshold of the q-value.

**Supplementary Figure 7.**
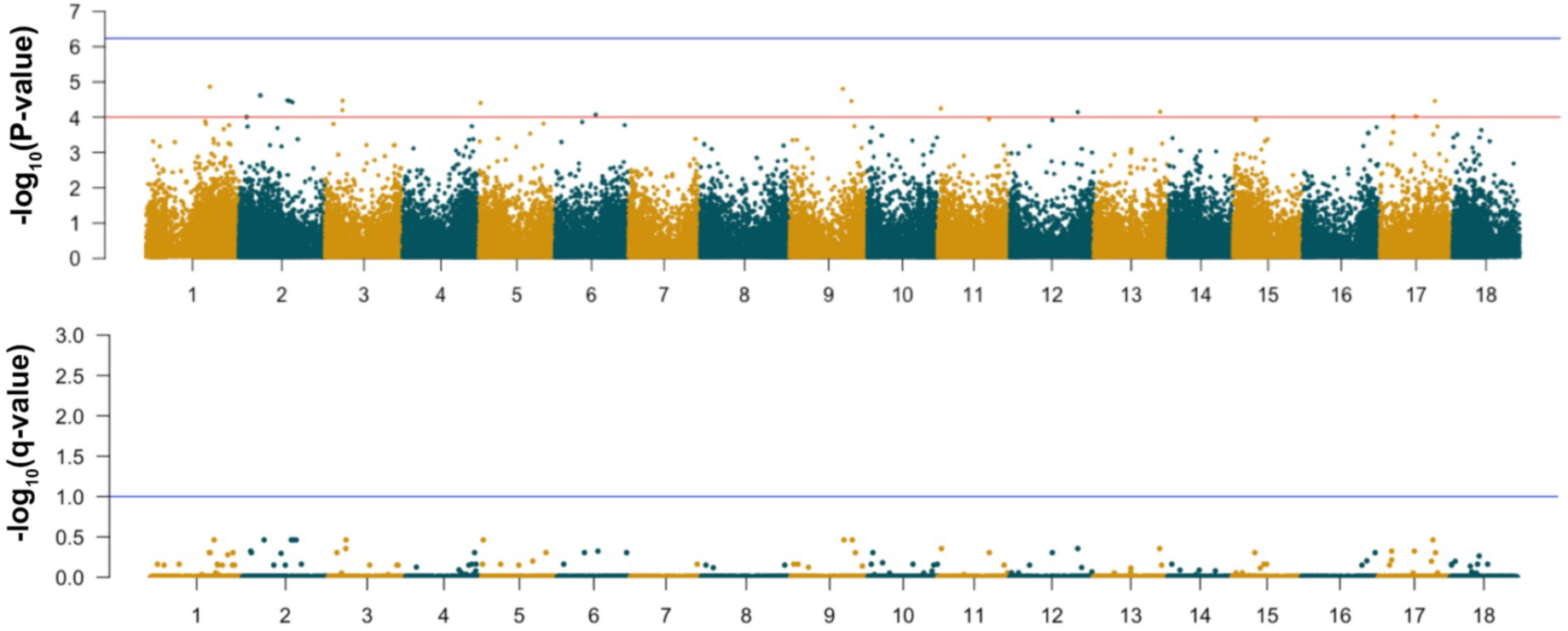
Multivariate GWAS of Area, Perimeter, Round, Solidity and AR (model 3). Manhattan plots of −log_10_(P-value) (top panel) and −log_10_(q-value) (bottom panel). In the top panel the blue horizontal line indicates the Bonferroni statistical threshold and the red line indicate a −log_10_(p-value) = 4. In the bottom panel de the blue line indicates the significant threshold of the q-value.

**Supplementary Figure 8.**
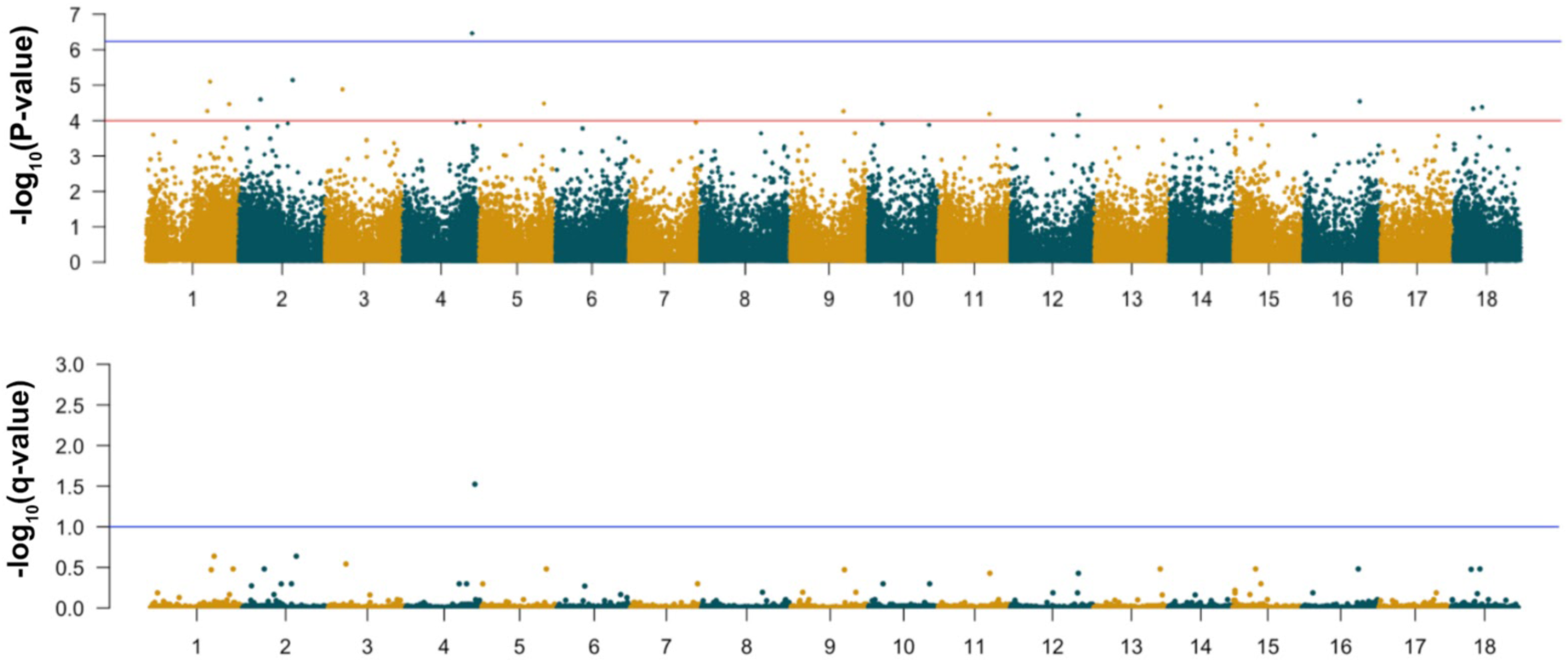
Multivariate GWAS of Area, Perimeter, Feret, Circularity, Round, Solidity and AR (Model 4). Manhattan plots of −log_10_(P-value) (top panel) and −log_10_(q-value) (bottom panel). In the top panel the blue horizontal line indicates the Bonferroni statistical threshold and the red line indicate a −log_10_(p-value) = 4. In the bottom panel de the blue line indicates the significant threshold of the q-value.

**Supplementary Figure 9.**
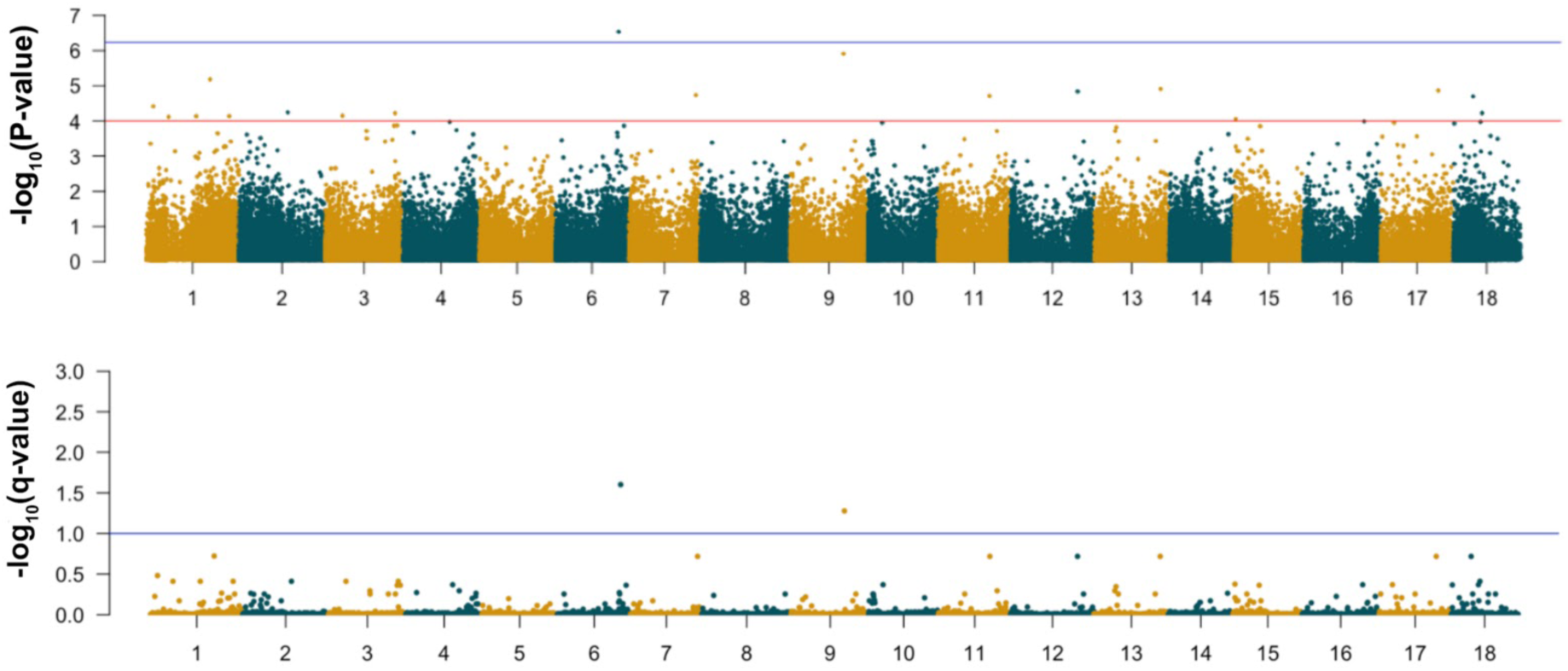
Multivariate GWAS of Area, Perimeter and Feret (model 5). Manhattan plots of −log_10_(P-value) (top panel) and −log_10_(q-value) (bottom panel). In the top panel the blue horizontal line indicates the Bonferroni statistical threshold and the red line indicate a −log_10_(p-value) = 4. In the bottom panel de the blue line indicates the significant threshold of the q-value.

**Supplementary Figure 10.**
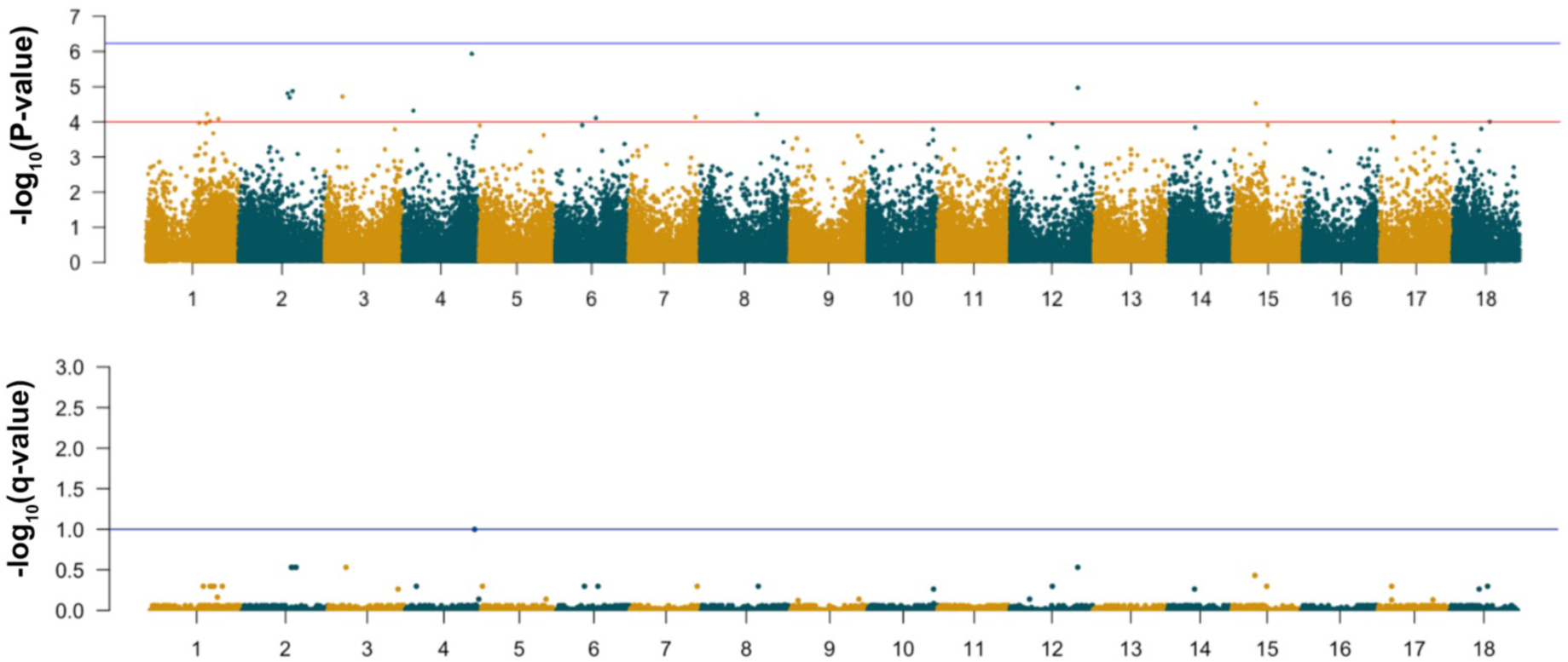
Multivariate GWAS of Circularity, Round, Solidity and AR (model 6). Manhattan plots of −log_10_(P-value) (top panel) and −log_10_(q-value) (bottom panel). In the top panel the blue horizontal line indicates the Bonferroni statistical threshold and the red line indicate a −log_10_(p-value) = 4. In the bottom panel de the blue line indicates the significant threshold of the q-value.

## Supplementary tables

**Supplementary table 1**. Mean values of raw and plot shape and size root measurements. The plot value was calculated as the average of five plants per plot, the raw value was the individual root value per genotype

**Supplementary table 2**. GWAS results of image-extracted and yield-related traits. Results in bold are SNP p-values that surpassed the Bonferroni threshold.

**Supplementary table 3**. GWAS results of image-extracted and yield-related traits using CMD adjusted phenotypes. Results in bold are SNP p-values that surpassed the Bonferroni threshold.

**Supplementary table 4**. GWAS results of the standard deviation of image-extracted traits using CMD adjusted phenotypes. Results in bold are SNP p-values that surpassed the Bonferroni threshold.

**Supplementary table 5**. Summary results genomic prediction. Input phenotypes: mean values per genotype/plot, mean adjusted phenotypes (mean + CMD correction), standard deviation per genotype/plot of adjusted phenotypes (sd + CMD correction).

**Supplementary table 6**. Candidate gene annotation of significant QTL regions in chromosomes 6,9 and 16 using the GWAS results of the standard deviation + CMD correction of root size and shape traits.

